# Label-free 3-D molecular imaging of living tissues using Raman Spectral Projection Tomography

**DOI:** 10.1101/2023.09.23.559025

**Authors:** Elzbieta Stepula, Anders R. Walther, Dev Mehrotra, Magnus Jensen, Mu H. Yuan, Simon V. Pedersen, Eileen Gentleman, Michael B. Albro, Martin A. B. Hedegaard, Mads S. Bergholt

## Abstract

The ability to image tissues in three-dimensions (3-D) with label-free molecular contrast at mesoscale would be a valuable capability in biology and biomedicine. Here, we introduce Raman spectral projection tomography (RSPT) for volumetric molecular imaging with sub-millimeter spatial resolution. We have developed a RSPT imaging instrument capable of providing 3-D molecular contrast in transparent and semi-transparent samples. A computational pipeline for multivariate reconstruction was established to extract label-free spatial molecular information from Raman projection data. We demonstrate imaging and visualization of phantoms of various complex shapes with label-free molecular contrast. Finally, we apply RSPT as a novel tool for imaging of molecular gradients and extracellular matrix heterogeneities in fixed and live tissue-engineered constructs and explanted native tissues. RSPT imaging opens new possibilities for label-free molecular monitoring of tissues.

## Introduction

Label-free imaging of tissues at mesoscale in three dimensions is of substantial importance across biomedical sciences. Mesoscale volumetric imaging techniques, characterized by intermediate spatial resolutions spanning from hundreds of micrometers to millimeters provide a comprehensive view of complex biological systems, allowing researchers to investigate relationships between cellular structures, tissue organization, and organ function. This comprehensive insight into the intermediate scale is essential for deciphering the multi-scale nature of tissues across different levels of organization.

Numerous three-dimensional (3-D) imaging techniques can be applied for imaging at mesoscale. Computed tomography (CT) and micro-CT uses X-ray absorption while magnetic resonance imaging (MRI) employs strong magnetic fields and radio waves to generate 3-D volumetric images.^1,2^ Optical methods for mesoscale imaging include optical projection tomography (OPT)^3^, optoacoustic tomography^4^, and diffuse optical tomography (DOT)^5^. OPT is used widely in biomedical research including in developmental biology, cancer research, and neurosciences^6^. OPT measures optical projections of the emission/absorption of light to generate volumetric images of transparent biological samples.^7^ All these techniques can provide volumetric spatial information at various scales; however, they do not offer direct label-free molecular contrast of the composition of the specimen.

Imaging at micrometer scale heavily relies on techniques such as confocal laser-scanning microscopy and light sheet microscopy that enables high-resolution volumetric imaging of labelled structures such live organoids or zebrafish models etc.^8–10^ Multiphoton imaging approaches such as second harmonic generation (SHG), two-photon excited fluorescence (TPEF), coherent anti-stokes Raman spectroscopy (CARS) and stimulated Raman spectroscopy (SRS) offers rapid label-free volumetric molecular imaging, but these have a small field of view (FOV) and are mainly limited to depths within a few hundred microns due to requirement of high numerical aperture objectives.^11–15^

Confocal Raman microspectroscopy can provide 3-D images of the chemical composition with diffraction limited resolution and have been used for imaging cells and organoids. Raman microspectroscopy is, however, limited in penetration depth to submillimetre size samples.^16–19^ Raman spectroscopy probes the specific molecular vibrations of the molecules offering information about the structure and composition of biological tissues. For molecular analysis in deeper tissues (from a few microns to several millimetres or centimetres) special techniques such as spatially offset Raman spectroscopy (SORS)^20–22^ have been developed. However, in the context of imaging, the major disadvantage of SORS is its limited spatial information due to the collection of diffusely scattered photons. For 3-D imaging, diffuse Raman tomography has been developed for non-invasive, label-free volumetric analysis.^23,24^ Diffusive approaches for image reconstruction require inverse modelling and are therefore combined with other techniques such as CT to obtain tissue geometry.^25,26^ Recent OPT approaches have captured isolated CH Raman bands using visible excitation.^27^ However, these techniques fall short in capturing the comprehensive molecular information present in full Raman spectra and are not compatible with applications involving living tissues.

Here we introduce Raman spectral projection tomography (RSPT) for volumetric label-free molecular spectral imaging at mesoscale. We have developed a transmission Raman spectral projection tomograph and established a comprehensive data analysis framework to extract molecular information from projected Raman spectra. We demonstrate accurate label-free 3-D molecular imaging of phantoms of various complex shapes supported by ray tracing simulations. We utilize RSPT as a new tool for evaluation of molecular gradients and spatial organization of extracellular (ECM) components in semi-transparent tissue-engineered (TE) constructs and native tissues. Finally, we demonstrate that the developed method can be applied on living tissues, highlighting the capability of the tool for temporal monitoring of TE cultures in their critical early phases of growth.

## Results

### Development of Raman spectral projection tomography

The development of Raman spectral projection tomographic imaging poses several inherent difficulties like those in OPT, along with additional prerequisites for image formation. Diffuse light is highly detrimental to any image formation in projection tomography. Laser excitation light and Raman scattered photons undergo multiple scattering events as they propagate through a tissue, resulting in breakdown of the parallel beam assumption with resulting blurring in the reconstructed image. Consequently, to form accurate and representative images, it is essential to capture the forward scattered Raman photons while rejecting diffuse light contributions from the sample. Secondly, traditional OPT captures full 2-D images at each angular projection while it would require new optical design to facilitate the acquisition of 2-D full spectrum Raman projections. Thirdly, the optical projections in a Raman spectroscopic setting would consist of compound Raman spectra accumulated through the sample, necessitating more sophisticated data preprocessing and analysis techniques to be developed.

We have developed a Raman spectral projection tomograph (RSPT) instrument based on transmission Raman geometry by rotation and translation of the sample (Figure 1A). In this design we excite and collect Raman photons from a single z-slice at a time. This has an additional advantage of reducing the collection of out-of-plane diffuse Raman photon contributions. The transmission Raman instrument provides benefits compared to a back-scattered (reflective) configuration, since transmission Raman allows for the accumulation of Raman signals emanating from deep within the sample, rather than Raman signals in proximity to the surface. We employed a custom manufactured high-power (2.0 W) near-infrared (NIR) 785 nm laser excitation allowing enough Raman signals to be generated within bulk ∼1 cm semi-transparent tissues. The incident laser beam was shaped into a line (power density of max ∼4 W/cm^2^) using a long focal length cylindrical lens and illuminated the sample similar to slice-illuminated optical projection tomography.^28^ The focal plane was situated at the midpoint of the rotation axis. As a result, each projection line-image contains Raman photons from the anterior part of the specimen (the portion closest to the laser excitation) alongside Raman signal contribution from the posterior back half of the specimen. To further filter the diffusely scattered light, we employ a long focal length and low NA imaging lens that predominantly collects the Raman scattered photons propagating parallel through the sample and project them onto a custom-manufactured line-to-line fiber bundle consisting of 43 cores (50 μm) acting as a slit. Inside the high-throughput spectrometer, the line-to-line fiber array was imaged onto a 2000×256-pixel charge coupled device (CCD) (*see materials and methods*). After binning the pixels representing individual fibers on the CCD, this configuration provides snapshots of 43 Raman spectra across the FOV. To accompany the instrument, we have developed a comprehensive controlling software for fully automated acquisition of data (Figure S1).

**Figure 1.**
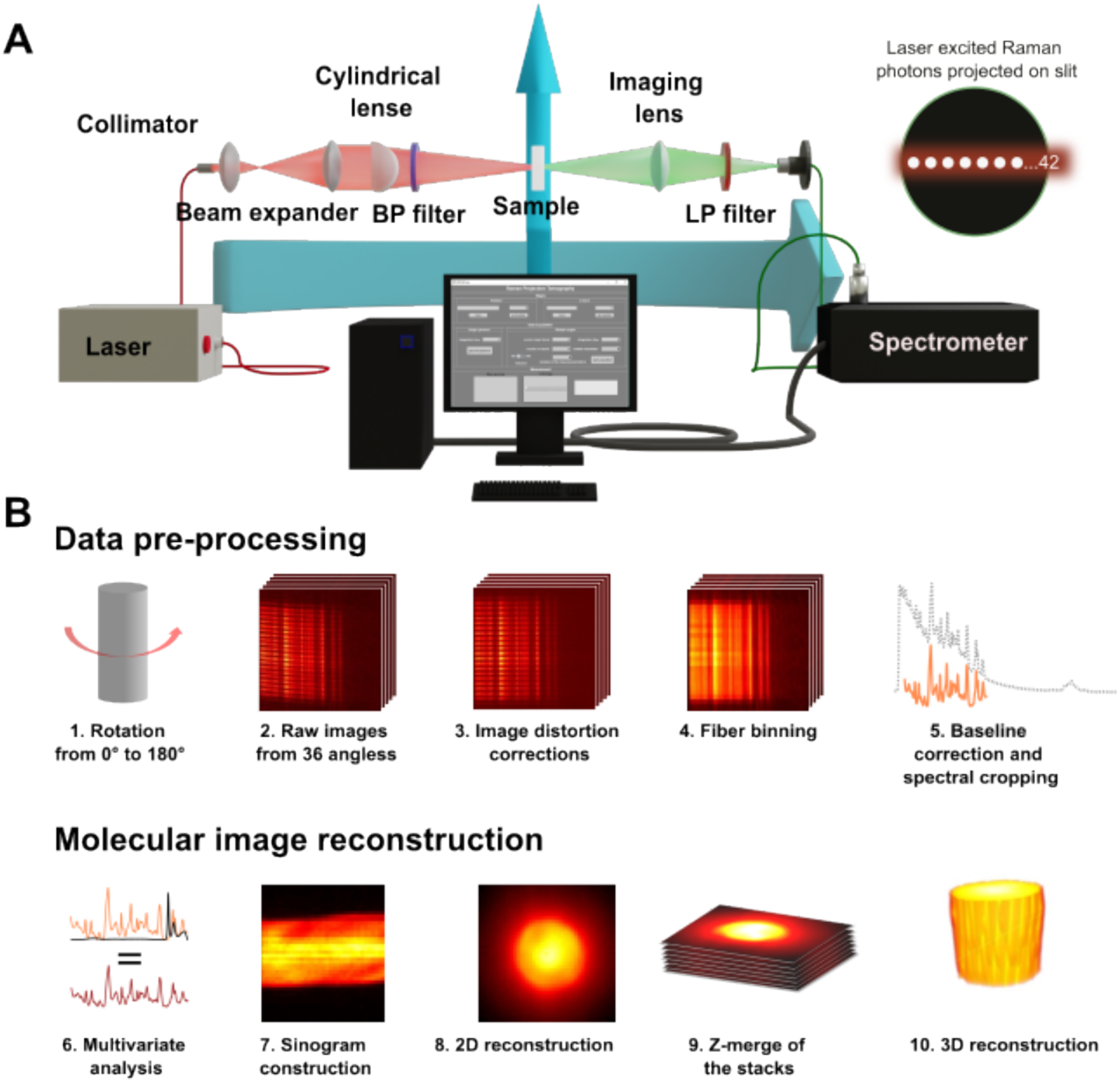
Raman Spectral Projection Tomography (RSPT) (A) Schematic illustration of the excitation and detection configuration for RSPT: A 2W 785 nm laser is beam shaped into a line using a cylindrical lens. The Raman scattered light is collected using a lens and imaged onto a line array comprised of 43 fiber cores coupled to a spectrometer. The sample is positioned for rotation around a 360° angle and z-axis displacement. (B) Depiction of data pre-processing and image reconstruction workflows: Pre-processing involves complex computational correction of image distortion in raw CCD images due to the spectrometer lens’s smile effect, fiber binning with mean calculations for 5 assigned pixel rows per fiber, and standard Raman processing including smoothing, baseline correction, and cropping to the spectral fingerprint region. The image reconstruction phase employs multivariate data analysis with pure component Raman spectra and 2-D reconstructions through inverse radon transformation. Sequential application to all z-slices enables the 3-D reconstruction.

Each projection plane generates a vast amount of hyperspectral Raman data (43 Raman spectra per projection angle), necessitating pre-processing, multivariate analysis, and reconstruction, which poses a significant computational challenge (*see Materials and Methods*). The developed data analysis framework incorporates data pre-processing (spectrometer image aberration correction, horizontal binning of fiber projections, wavelength calibration, autofluorescence subtraction and normalization). To enable molecular specific reconstruction, we employ multivariate regression of purified components.^29^ 3-D image reconstruction and rendering can finally be performed using inverse Radon transform - backprojection of the regression component abundances (Figure 1B).^30^

### Benchmarking of RSPT imaging performance

The developed projection system has a FOV of ∼9 mm (Figure 2A) with a magnification factor of the projection images on the fiber bundle of ∼0.4 and a voxel size of 0.2 x 0.2 x 0.1 mm after volumetric reconstruction. To characterise the system’s contrast, we fabricated three groups of spatial resolution targets with aperture widths of 1000, 500, and 400 µm, using transparent resin. These targets were imaged at a 45-degree angle to the optical axis (Figure 2B). This diagonal arrangement allowed us to assess the system’s capacity to resolve details at varying depths within the sample. From each sample, we collected Raman projections of the CH2 stretching vibration of the polymer (2940 cm^-1^) along the depth of field (Figure 2B).^31^ We then calculated the contrast ratio defined as the intensity difference between the resin target aperture and background (*see material and methods*).

**Figure 2.**
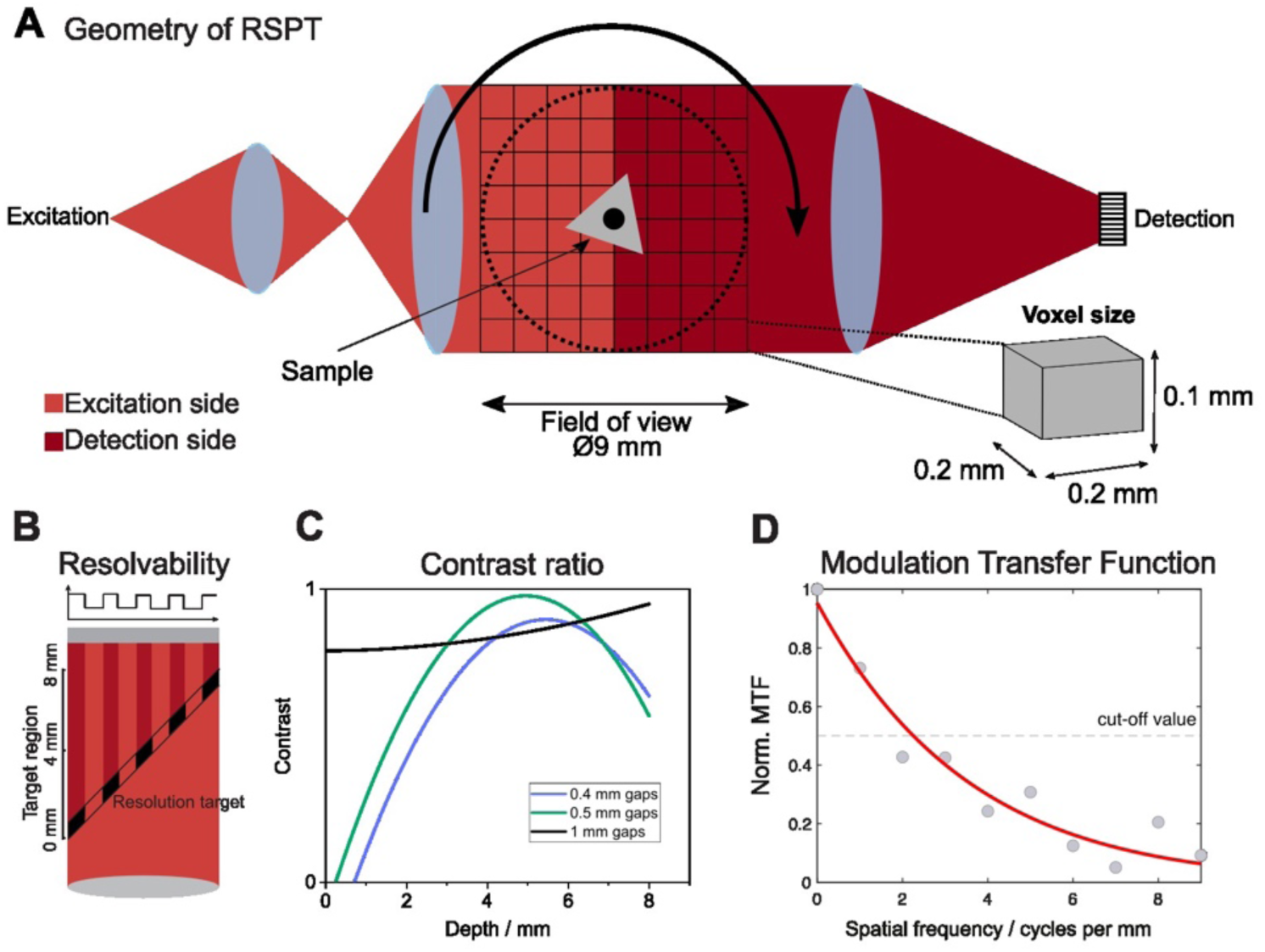
Benchmarking of Raman Spectral Projection Tomography. (A) Schematic representation of the RSPT setup, detailing the laser excitation (red) and Raman detection (carmine). The centrally located sample undergoes rotation within the 9 mm field of view, with an inset showcasing the experimentally defined voxel dimensions. (B) Evaluation of the RSPT’s spatial resolution using resolution targets that generates Raman signals. These 3-D printed targets, made from resin, were structured with three distinct aperture widths. The target’s midpoint was positioned at the cylindrical lens’s focus, with a 45° incline relative to the optical system’s perpendicular axis. (C) Resultant measurements facilitated the derivation of a plot indicating the relationship between the contrast ratio and the depth of field in RSPT imaging. (D) The calculated MTF for the RSPT system were based on Edge Spread Function from a 2-D reconstruction of a cylindrical sample. From this, the Line Spread Function (LSF) was derived and the MTF was computed by determining the magnitude of the Fourier Transform of the LSF. A demarcated cut-off at the 50% MTF threshold reveals the system’s empirical resolution capability.

To quantify the RSPT system’s overall performance, we calculated the modulation transfer function (MTF) (Figure 2B). This provides a quantitative assessment of the spatial resolution and ability to reproduce varying levels of detail from the object onto the image plane (see *Materials and methods*). The MTF illustrates the system’s performance across a range of spatial frequencies. By examining specific points on the curve, such as the frequency at which the MTF drops to 0.5, we can determine the system’s effective resolution limit to 600 µm. Considering our voxel size of 0.2 x 0.2 x 0.1 mm, the system meets the Nyquist sampling theorem for our determined resolution limit, ensuring that spatial details are captured and undersampling artifacts are minimized.^32^

### Label-free 3-D imaging of phantoms with molecular contrast

Supported by ray tracing simulations (Figure 3A), we evaluated the capability of RSPT imaging to reproduce 3-D printed semi-transparent resin samples with varying complex shapes (cylinder, cuboid, triangular prism and a combination of cylinder and cuboid) (Figure 3B). A pure reference spectrum of resin (Figure 3C) was measured for multivariate regression analysis. We calculated the sinograms showing all projections of a plane (0° to 360°) and 3-D rendering of the molecular abundance of resin. The shapes of the 2-D and 3-D reconstructions largely replicate the structures of the phantom samples and simulations (Figure 3D). This data indicates that RSPT can provide reliable 3-D molecular imaging of complex transparent shapes.

**Figure 3.**
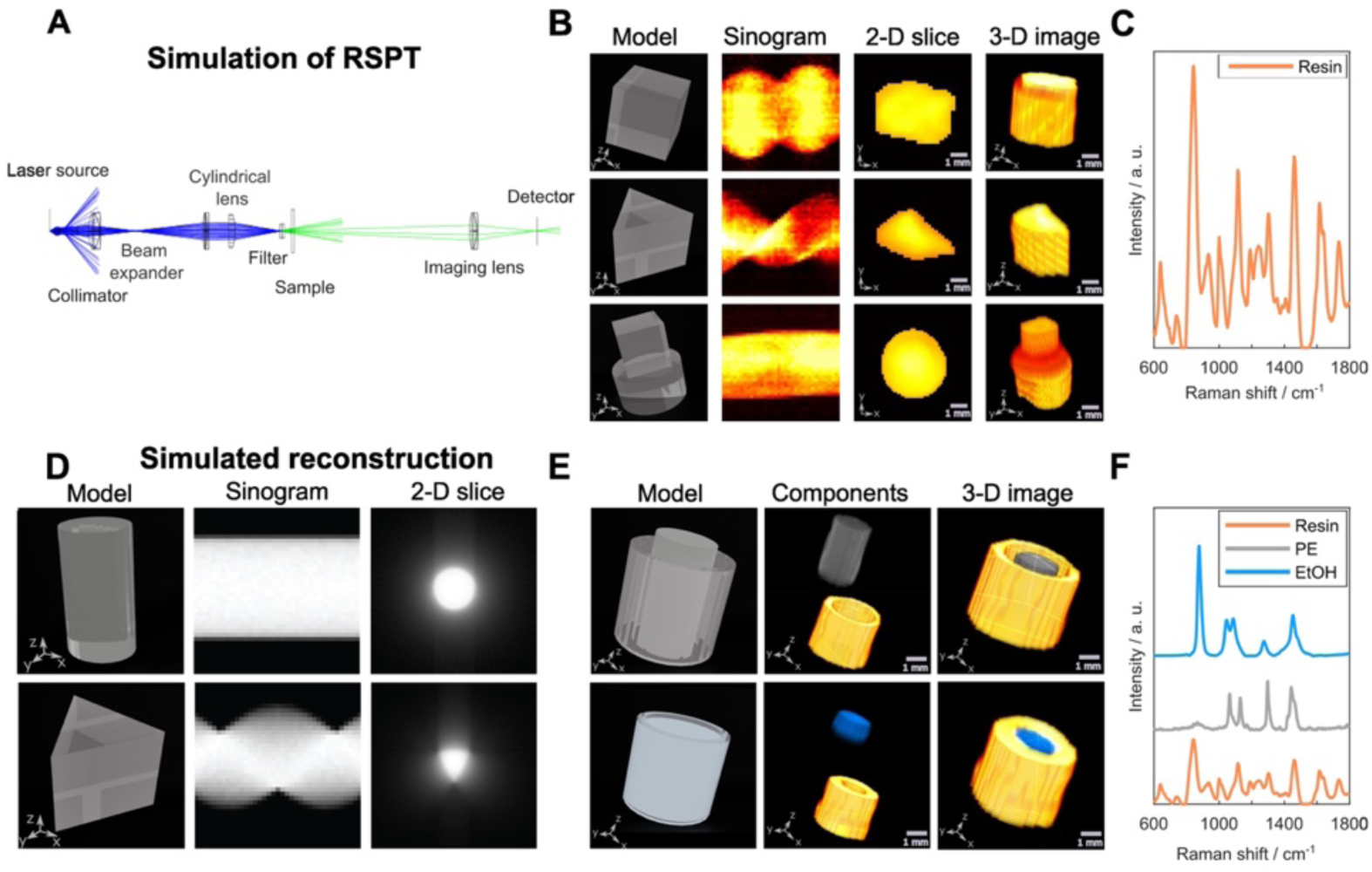
Simulated and experimental results for phantom RSPT imaging. (A) Schematic representation of the RSPT prototype simulated in Zemax OpticStudio. Ray tracing demonstrates the predominant detection of forward-scattered Raman photons in the designed system. (B) Comprehensive visualizations of 3-D printed single-component phantom samples using resin, with diverse geometries: cuboid, prism, and an amalgamation of cuboid and cylinder. The 2-D reconstructions showcase the basal slice of each sample, whereas the 3-D renditions encapsulate measurements spanning 10 slices. (C) Pure reference Raman component of resin. (D) Depiction of sinograms and 2-D reconstructions derived from Zemax-simulated ray traces of cylinder and prism-shaped samples. (E) 3-D representations of two component samples: a hollow resin cylinder encasing a polyethylene (PE) rod and the inferior filled with a pristine ethanol solution. Subsequent columns show the 3-D reconstructions of individual components based on their Raman molecular signatures. (F) Measured reference Raman spectra of the constituent components.

We then evaluated the capability of RSPT imaging to reconstruct hollow cylindrical-shaped semi-transparent resin phantoms with two profoundly different molecular components. We imaged a resin cylinder and an opaque polyethylene (PE) rod with a smaller diameter placed in the center as well as a cylinder filled with ethanol (EtOH) (Figure 3E). 3-D reconstructions of the phantoms based on the Raman spectra of pure components (Figure 3F) demonstrate an exceptional capability to create a label-free molecular contrast at mesoscopic scale.

### RSPT imaging of ECM in native and living tissue engineered constructs

We next investigated if RSPT imaging could be used for label-free bioimaging of live and devitalized tissue specimens, albeit with lower spatial resolution than phantoms due to optical scattering and therefore partial breakdown of the parallel ray assumption. Tissue engineering (TE) is a growing therapy degenerative pathologies (e.g. musculoskeletal, cardiovascular) that aims to regenerate functional tissues for clinical replacement of damaged or diseased tissue. TE platforms aims to adopt regenerative protocols using a variety of biomaterials, cells, and growth factors to generate repair tissues that recapitulate the composition, structure, and function of native tissue.^32–34^ Given the influence of ECM organization on tissue functionality (e.g. mechanical load transmission),^33–35^ the ability to achieve non-destructive, label-free quantitative imaging of the spatial distribution of ECM may be critical for the clinical evaluation of the quality of engineered tissue grafts.^36,37^ For cartilage TE, protocols aims to regenerate neocartilage with a native-matched cartilaginous ECM, comprised of high levels of glycosaminoglycans (GAGs) interspersed within a type II collagen fibril network, which give rise to the tissue’s unique ability to provide lubricity and mechanical load support.

TE cartilage constructs were fabricated via seeding of bovine chondrocyte in agarose scaffolds and cultivated for 42 days. RPT endpoint imaging was performed on constructs at day 0 or day 42, as well as on native bovine cartilage explanted tissues for reference, to evaluate ECM heterogeneities (Figure 4A). The Raman spectra of pure reference molecules (GAG, type II collagen, agarose, and water) were measured for the multivariate regression analysis and 3-D reconstruction (Figure 4B). Molecular heterogeneities were extracted from the principal x-z slices (Figure 4C). RSPT imaging of the TE constructs at day 0 showed as expected a background signal of low intensity scores of collagen and GAG largely indicating the absence of these components. By day 42, chondrocytes had deposited significant levels of GAG and collagen. ECM was distributed in a spatially dependent manner with constituents concentrated at the culture-media-exposed construct periphery (top-most region) but more dilute in the tissue center (bottom-most region), consistent with prior *ex vivo* characterizations of ECM heterogeneities that result from nutrient/growth factor transport gradients in TE applications.^37–38^ Conversely, native cartilage explants exhibited inverse heterogeneities with GAG and collagen constituents concentrated in the lower tissue regions, consistent with their established spatial distribution through the depth-dependent zonal regions in native cartilage.

**Figure 4.**
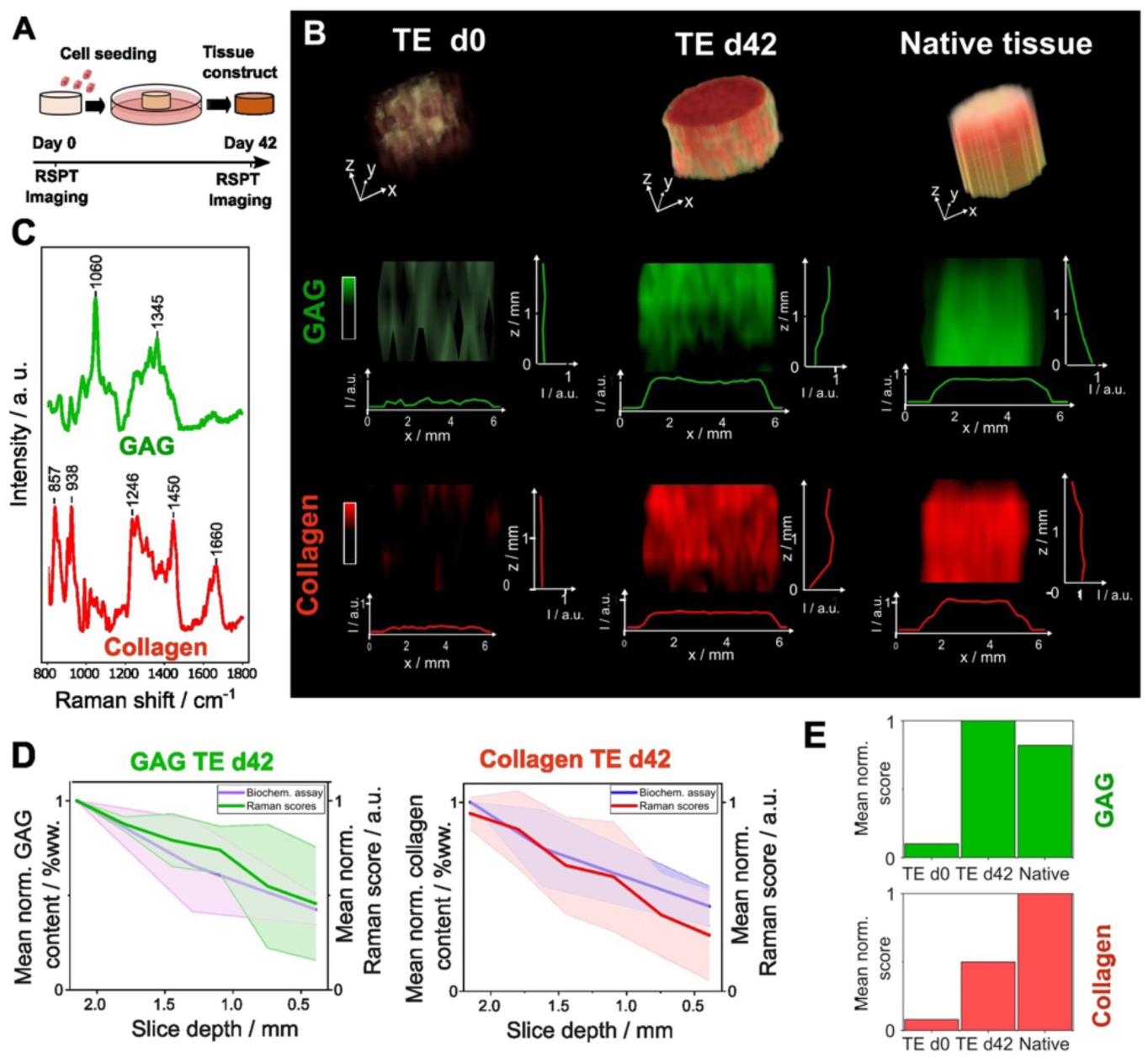
RSPT imaging of tissue-engineered constructs vs native tissues. (A) Schematic of the culturing of the tissue engineering constructs, where chondrocytes were seeded in agarose and cultured in chondrogenic media for 42 days. (B) The uppermost row shows 3-D rending of the total GAG + collagen for day 0, day 42 TE cartilage, and native cartilage. Also shown are virtual x-z slices revealing the molecular gradients within each specimen. (C) Normalized Raman spectra of references (collagen and GAG) employed in multivariate data analysis across native cartilage and tissue-engineered (TE) cartilage at day 0 (d0) and day 42 (d42). (D) Comparison of the results of biochemical assays assessing the %ww content of both collagen and GAG in three zones of the day 42 TE constructs samples with the mean Raman scores for each slice measured with RSPT. (E) Normalized mean sum of GAG (green) and collagen (red) contents for d0, d42, and native samples, expressed as a proportion of the total sum of all components (GAG + collagen + water). While GAG content is minimal at d0, it experiences a significant elevation by d42, surpassing that in native samples. In contrast, collagen content at d42 is reduced in comparison to its presence in native samples.

Correlation analysis of the molecular gradients with biochemical assays (1,9-dimethylmethylene blue (DMMB) assay for GAG and Hydroxyproline assay for collagen) demonstrated excellent agreement (Figure 4D). Collagen and GAG concentrations detected by RSPT imaging were highest at the surface and decreased deeper into the TE constructs. For native tissues the gradients were opposite and increased deeper in the tissues in agreement with previous reports using microscopy and histology^37–38^. Additionally, we semi-quantitatively assessed the composition of GAG and collagen (Figure 4E) showing a significantly increase in GAG by d42 even surpassed that observed in native samples. Contrarily, the collagen content demonstrated a gradual rise towards the native levels. Hence, RSPT imaging represent a very powerful tool for non-destructive bulk tissue gradient and heterogeneity molecular analytics. It is important to note that the RSPT imaging indicated notably weak signals for water and agarose components. This suggests that RSPT may not be the most optimal method for discerning the concentrations of these two components. In a parallel tissue culturing experiment (*see Material and Methods*), we investigated whether tissues could be imaged in a live setting. Live chondrocytes were encapsulated in alginate hydrogels and divided into an exposed group and a control group. The exposed group was subject to imaging with the RSPT system (duration ∼3 hours), the control group was taken out of the petri dish for the same duration and was subject to the same hydration as the measured sample. Afterwards a Live/Dead Cell Viability Assay was performed on both samples. The staining revealed that upon exposure to 2W laser power had negligible effects on cell viability (Figure S3). The percentage of live cells in the laser-exposed sample was 73%, while the control group exhibited a slightly lower cell viability at 62%.

## Discussion

At present, there are limited tools for mesoscale imaging that enable direct visualization of the molecular makeup of the ECM without requiring labelling. Raman spectral projection-based tomography offers volumetric imaging of bulk samples and effectively complements traditional Raman microscopy, such as its inability to capture depth information, as well as those of SORS, which offers depth dependent data but lacks imaging functionalities.

There are two main mechanisms enabling image formation in the developed technique. The instrument uses slice illumination, and the projections of Raman spectra are acquired using a one-dimensional fiber array. This effectively rejects Raman photons scattered out of the imaging plane reducing lateral blurring. Secondly, although the long working distance imaging lens results in lower effective numerical aperture and less signal being detected it ensures that mostly forward scattered Raman photons are captured. Hence, RSPT employs a similar approach to OPT where the imaging lens acts as a collimator, reducing the collection of diffuse photons.

We demonstrated the capability of RSPT imaging to volumetrically replicate the geometry of resin phantoms with high molecular contrast. The minor artifacts introduced during the image generation process can largely be attributed to refractive index mismatch. Immersing real tissue samples in refractive index matching liquids to eliminate refractive mismatches might be feasible but would require a sizeable Raman transparent cuvette. The reconstruction of phantoms with different molecular components demonstrates that the compound Raman spectra can largely be deconvolved using multivariate regression analysis coupled with backprojection of the molecular abundances.

We applied the technique in a TE setting aiming to produce functional tissue replacements. Mimicking the native cartilage’s ECM content and structural organization, a determinant of implant longevity, remains challenging.^38^ Hence, the availability of a new semi-quantitative and non-destructive imaging tools such as RSPT is vital for temporally assessing living engineered tissues. Projection tomography relies on the assumption of optical transparency in the samples under imaging. However, when samples scatter light, it leads to a notable decrease in both image resolution and quality. The constructs and native tissues used represent ideal samples consisting of collagen, GAG, agarose, and water. The agarose scaffold is optically transparent, and the absence of vascularity and light absorption enables relatively intense Raman signals to be generated and captured even through ∼8 mm of bulk TE and native tissues. The imaging results largely mirrored the ECM heterogeneities of engineered and native cartilage specimens and aligns with previous microscopic findings using confocal Raman microscopy of tissue sections as well as biochemical assays.^39^ Moreover, the system proficiently imaged live TE constructs comprising chondrocytes in hydrogel matrices indicating that prolonged exposure (<180 min) to a 2.0 W 785 nm laser excitation does not compromise cell viability. This is likely due to the efficient heat dissipation and absence of any significant chromophores in the sample.

RSPT imaging comes with limitations. The sample size is restricted due to the limited penetration capability of laser light in semi-transparent tissues. The reconstruction will therefore largely be sample dependent and requires prior knowledge of the key molecular makeup. Further the computational framework assumes a linear relationship between molecular abundance and Raman signal intensity. Since the laser excitation has a gaussian profile and diffuses in tissue any analysis of the absolute Raman signal intensity remains difficult. Normalization of the compound Raman spectra projections are therefore of paramount importance in obtaining relative gradients.

There is considerable potential for further enhancing RSPT. Here we used simple backpropagation reconstruction but more advanced reconstruction methods such as iterative reconstruction algorithms and scattering correction factors could potentially enhance the 3-D rendering^40^. Further, there are multiple avenues to explore for improving instrumentation including incorporation of a telecentric optic design, slice illumination in both x, y- and z, direct imaging on to spectrometer input slit or even wavefront shaping approaches for better laser excitation control. Exploring the integration of multimodal OPT and RSPT presents exciting prospects for delving into functional imaging using diverse contrast mechanisms.

## Conclusion

Raman spectral projection tomography (RSPT) offers label-free volumetric molecular mesoscale imaging of transparent and semi-transparent samples with sub-millimetre spatial resolution. The molecular reconstruction enabled accurate replication of transparent phantoms. The RSPT imaging technique could generate 3-D molecular contrast and enabled the temporal semi-quantification of molecular gradients in bulk tissue. These findings show that the imaging approach can serve as a platform for minimally invasive molecular analysis and a means to meticulously track molecular gradients within living tissues with high precision in both contrast and resolution.

## Material and Methods

### Raman spectral projection tomography instrumentation

The Raman spectral projection tomography instrument (Figure 1A) consists of a rotary stage (HDR50/M Thorlabs) to provide a full 360° rotation of the sample. A custom wavelength stabilized 785 nm laser with maximum output power of 2.0 W (Innovative Photonic Solutions) is fiber-coupled through a 365 µm low-OH multi-mode fiber and collimated using an aspheric condenser lens and then expanded using a Keplerian lens system (*f* = 30 and *f* = 100 mm, Thorlabs). To obtain light sheet illumination in the sample plane we used a plano-convex round cylindrical lens (*f* = 50 mm, Thorlabs). The laser light is finally passed through two 785 nm MaxLine laser clean-up filters (LL01-785-25, Semrock) to remove optics background signals. Samples were placed on a motorized linear translation stage (LTS300/M Thorlabs) allowing movement in the z-direction. The generated Raman signals along the sample is imaged directly onto a custom-made linear fiber array (43 fibers, 50 µm) (Ibsen Photonics A/S) acting as a spectrometer slit using an imaging lens (*f* = 50 mm, Thorlabs). A set of three 785 nm EdgeBasic long-pass edge filters (BLP01-785 R-25, Semrock) was used on the detection side to remove the Rayleigh scattering signal from the intense laser light. The linear fiber bundle is directly coupled into a high-throughput Raman spectrometer (Eagle, Ibsen Photonics) equipped with a back-illuminated deep-depletion CCD (Andor iVac, Oxford Instruments). Since tissue samples are susceptible to dehydration under the influence of high-intensity laser exposure we integrated an ultrasonic humidifier to avoid the samples from drying out.

### Instrument control software

We developed software for controlling our RSPT instrument in MATLAB (version 2020b, MathWorks Inc.), including a graphical user interface (GUI). The GUI was designed to record single projections as well as a complete sequence of projections, where the integration time of the measurement, angle rotation resolution and z-distance between the slices can be set. Additionally, a raw CCD image and mean spectrum of the ongoing measurement is displayed for real-time visualization of the data quality during acquisition.

### Raman image processing and 3-D molecular image reconstruction

All spectral processes and analyses were performed within the MATLAB environment (version 2020b, MathWorks Inc.) (Figure 1B). Initially, the spectrum underwent wavelength calibration via a Mercury-Argon source (Ocean Insight). Subsequent steps were used to correct aberrations of the spectrometer. For the aberration correction a broadband light source spectrum was captured, providing registration points for polynomial transformation. The derived spatial transformation was then autonomously applied to all the projection datasets. The following process involved spatial software binning for the 43 fibers present on the CCD camera, with each fiber comprising 5 rows of pixels. Prior to the next steps, the spectra were cropped to the fingerprint wavenumber range spanning 800-1800 cm^-1^. The cropped spectra were then smoothed utilizing a Gaussian filter with σ=2.^41^ Subsequently, a 3^rd^ order constrained polynomial fit for baseline correction was employed for effective elimination of autofluorescence. It should be noted that as the lasers excitation profile is Gaussian-like, normalization of Raman spectra to their integrated areas is key to enable semi-quantitative molecular analysis.

Molecular contrast from the tomographic imaging data set is derived by applying nonnegative constrained least-squares (NNLS) method. Based on the reference spectra using NNLS we calculate the abundance of each compound. Sinograms for each of the compounds were constructed by combining false color abundance images for every measurement angle in one figure, resulting in one sinogram per every pure component. A standard backprojection (BP) algorithm was applied for the image reconstruction from each component sinogram. To ensure consistent value scaling across the values of the scores, min-max normalization was applied, transforming all feature values to lie within the 0-1 range. The precise workflow for RSPT image reconstruction including the CCD images and spectra is presented in Figure S4.

The gradient trends for each component are shown by calculating the mean score of the component for every slice (excluding background) across and along the sample axis and plotting it against the slice number which corresponds to the sample zone.

### Benchmarking of Raman Spectral Projection Tomography instrument

The FOV is limited by the imaging lens, linear detection fiber array as well as the width of the line-shaped laser beam. The dimensions of the FOV in the RSPT instrument are 9 mm x 9 mm. The voxel size can be calculating by ratio of the FOV to the number of detectors (43). The magnification was calculated by dividing the size of the image on the detector by the actual size of the object.^42,43^

The contrast ratio of RSPT as a function along the optical axis was measured using a set of molecularly sensitive resolution targets that were 3-D printed (Zortrax, Inkspire). These targets have different sizes of line patterns (1 mm, 0.5 mm, and 0.4 mm). They were positioned at a 45-degree angle to the optical axis and three projections with an integration time of 30 s were captured. The contrast ratio was derived from the intensity variations between the peaks and valleys of the subtracted data. This ratio provides a quantifiable measure of the system’s resolution prowess. An additional second-degree polynomial fit extended our understanding of contrast dynamics across the sample, underscoring the system’s capability in detailed feature differentiation.

The Modulation Transfer Function (MTF) was ascertained using a 2-D reconstruction of a cylindrical sample. We opted to assess the MTF on a reconstructed sample to achieve a realistic representation of our system’s performance under typical imaging conditions. This approach not only accounts for possible reconstruction artifacts but also offers a basis for meaningful comparisons with other imaging systems. Within the reconstructed image, a region of interest (ROI) with a pronounced contrast edge was pinpointed. The Edge Spread Function (ESF) was then determined by averaging pixel values in the ROI along the edge direction. From this, the Line Spread Function (LSF) was derived by differentiating the ESF. The MTF was subsequently computed by evaluating the Fourier Transform of the LSF and determining its magnitude. To ensure normalization, this MTF was divided by its peak value and then graphed against spatial frequency, providing an insight into the spatial resolution and overall image quality. A 5^th^ order polynomial fitting was employed to represent the MTF, as showcased in Figure 2C, where the MTF was charted versus the spatial frequency (cycles per mm).

### Optical simulations

The RSPT instrument was simulated using Zemax OpticStudio 14.2 software. The simulation included two bulk samples with distinct geometries: a cylindrical and a triangular sample. To approximate scattering, we employed Mie scattering.^44^ The parameters used for Mie scattering, such as mean path, particle index (refractive index), size, density, minimum threshold, and transmission, were estimated based on the optical properties of 3-D printed resin.^45^ The proportion or fraction of incident light that passes through a material or sample, the transmission parameter was estimated using Lambert-Beer’s law and the absorption coefficient.^46^ Additionally, we developed a custom script that virtually rotates the sample and records the stimulated pixel intensity at the detector for each rotation. To process the simulation results for 3-D reconstruction, we employed a custom MATLAB script that imports the detector data, generates a sinogram, and reconstructs the final image. This approach allowed us to successfully model and analyze the performance of the RSPT system.

### Phantom samples development and imaging

Single component phantom samples in various shapes (cylinder, cuboid, triangular prism and a combination of cylinder and cuboid) were designed with Autodesk Inventor software and then 3-D printed with a UV resin printer (Zortrax, Inkspire) using semi-transparent resin. Two component samples include a resin hollow cylinder with an inserted polyethylene cylinder. The second two component samples include a resin hollow cylinder filled with isopropanol (PA, Merck). We acquired a full 360° scan with projections taken every 20° with an integration time of 10 seconds, the z-step size was 500 µm and a total number of 5 slices was recorded.

### Native cartilage tissue

Articular cartilage explants were obtained from the medial and lateral condyles of 3–6-month-old bovine calves (Green Village Packing Co., Green Village, NJ) using a 04 mm biopsy punch. Explants were then cut at a depth of 3±0.1 mm from the superficial zone surface, removing the subchondral bone. Afterward, specimens were preserved utilizing a 4% formaldehyde solution in deionized water, supplemented with ethanol and acetic acid, for a duration of 24 hours. After the fixation process, the samples underwent three successive 1x phosphate-buffered saline (PBS) washes, with each wash lasting 24 hours. Following this protocol, the specimens were securely stored in PBS buffer.

### Culturing of tissue engineered constructs

Primary articular chondrocytes were isolated from eight bovine carpometacarpal joints (3-6 months old, Green Village Packing Co). Harvested cartilage was digested using Type IV Collagenase (Worthington) overnight at 37°C for 17 hours with orbital shaking. Isolated chondrocytes were seeded in 2% w/v agarose (type VII, Sigma) at 30 × 106 cells/mL and cast into 2 mm-thick molds constructed using glass slides with a rubber spacer. Cylindrical tissue constructs were procured via a Ø6 mm biopsy punch. Tissue constructs were cultured in chondrogenic media consisting of high glucose Dulbecco’s Modified Eagle’s Media (DMEM, Gibco) supplemented with 1 mM sodium pyruvate, 50 µg/mL L-proline, 100 nM dexamethasone, 1% ITS+ premix (Corning), 1% PS/AM antibiotic-antimycotic, and 50 µg/mL ascorbate-2-phosphate (Sigma) at 37 °C with regular media changes (three times per week) for a duration of 42 days. Tissue constructs were cultured in the presence of 10ng/ml TGF-β3 (human recombinant active TGF-β3, R&D Systems) for the initial 14 days, after which TGF-β3 supplementation in the media ceased. Samples were removed from culture for tissue characterizations at days 0 and 42 and fixed in a solution containing 4% formaldehyde in deionized water, complemented with ethanol and acetic acid, for an uninterrupted 24-hour period. After the fixation, the samples were methodically rinsed in 1x phosphate-buffered saline (PBS) thrice, with each rinse spanning 24 hours. Post-rinsing, the specimens were stored in a PBS buffer.

### Glycosaminoglycan (GAG) and collagen assays

The quantification of glycosaminoglycan (GAG) was conducted using an assay rooted in the metachromatic interaction between sulfated GAGs and a designated dye. This binding interaction leads to an observable shift in the absorption maximum, rendering the solution’s color to change from blue to a pronounced purple in response to increasing GAG concentrations. To standardize the assay, a series of standards, ranging from 0 to 50 μg/mL, were prepared. Subsequent sample digest dilutions were introduced to the respective wells, followed by an equal volume of the specific dye. Post-incubation, the absorbance values of the samples were measured and juxtaposed against those of the known standards. GAG concentrations in the samples were then discerned, factoring in the dilution and sample digest volume. The outcomes were subsequently normalized to the tissue weight to express the GAG content in terms of (%ww).^47^

The quantification of collagen hinges on the presence of the amino acid hydroxyproline in cartilage collagens. Given that hydroxyproline, along with proline, contributes to roughly a quarter of the amino acid content in collagen, its quantitation serves as an indirect measure of collagen content. For the preliminary step, each sample digest was treated with 12N HCl, facilitating the liberation of hydroxyproline from its polypeptide structure. The samples were then sealed in ampoules and incubated at 110°C. Following hydrolyzation, acid evaporation was achieved by heating the ampoules at 50°C until dryness. The resultant dried samples were reconstituted in hydroxyproline assay buffer and subjected to overnight refrigeration.The assay phase initiated with the oxidation of free hydroxyproline to pyrrole using chloramine-T. Subsequent introduction of p-Dimethylaminobenzaldehyde culminated in the formation of a chromophore, whose absorbance was measured spectrophotometrically. Using hydroxyproline standards with established concentrations, collagen content was extrapolated from the assay readings. Factoring in the assay buffer volume, the sample dilution ratio, and the known weight ratio of hydroxyproline to collagen, the collagen content was determined and normalized against the sample weight to derive the collagen concentration in terms of (%ww).^48^

### Encapsulation of chondrocytes in alginate hydrogels

Alginate hydrogels were fabricated as described previosuly.^49^ Briefly, sodium alginate (Sigma-Aldrich, 71238) was treated with 3 Mrad gamma irradiation to obtain low molecular weight alginate. Low molecular weight alginate was then dissolved and dialyzed (35 kDa cutoff) in milliQ water for three days. The final product was purified with activated charcoal, sterile filtered, frozen at -20°C and lyophilized. Chondrocytes were isolated from calf chondrocytes and cultured in the growth medium containing hgDMEM (Thermofisher, 42430082), 10% (v/v) fetal bovine serum, 1% (v/v) Antibiotic-Antimycotic (Thermofisher, 11570486) and 1% (v/v) nonessential amino acids (Thermofisher, 11140050). Cell encapsulation was performed as described previously.^50^ Confluent chondrocytes at passage one were dissociated and resuspended in serum-free DMEM at a final concentration of 50 million cells ml-1. 3% (w/v) alginate in in serum-free DMEM)and containing cells was placed in a Luerlock syringe (VWR, LOCA120006IM). 488mM calcium sulfate in serum-free DMEM was loaded into another Luerlock syringe and mixed with the cell-alginate solution through a Luerlock connector. The mixture was then deposited between hydrophobic glass plates spaced 1 mm apart and allowed to gel for 45 minutes. Cell-laden hydrogels were cut into smaller pieces and cultured in growth medium supplemented with 0.05 mg ml^-1^ L-ascorbic acid-2-phosphate (Sigma) and 1 mM calcium chloride (Sigma). Cell-laden were cut into 6x6 mm squares and cultured in growth medium supplemented with 0.05 mg ml-1 L-ascorbic acid-2-phosphate (Sigma) and 1 mM calcium chloride (Sigma) for 21 days.

### Viability assay

Live/Dead (Invitrogen, L3224) staining was performed according to manufacturer’s instruction. In brief, 500 µl cDPBS (containing 1 µM calcium chloride) with 2 µM Calcein AM, 4 µM ethidium homodimer-1 and 1 µg ml-1 Hoechst 33342 (Thermofisher, 62249) was added to chondrocyte-laden alginate hydrogels and was incubated for one hour. Hydrogels were then washed twice in DPBS containing 1µM calcium chloride and imaged on a laser scanning confocal microscope (Zeiss).Cell viability was assessed using ImageJ software. After segmenting and counting the stained cells in the acquired images (30 images per sample), the percentage of live cells was calculated by dividing the number of live cells by the total cell count (live plus dead cells).

### RSPT imaging of tissue engineered constructs and native cartilage

We performed 360° scan with projections taken every 20° with an integration time of 30 seconds per projection to accumulate enough Raman signal. Samples had diameters around 5 mm and a height of approximately ∼2 mm. We measured 7 slices, with a 300 µm resolution. The samples were hydrated intermittently with 5 µL of phosphate-buffered saline (PBS) between each slice measurement. We performed imaging with a 360° rotation resolution of 20°, and an integration time of 45 seconds. However, measuring tissue-engineered cartilage presented more challenges than native samples. Prior to the commencement of the RSPT assessment, a volume of 10 µL of Phosphate-Buffered Saline (PBS) were introduced into the sample holder. This procedure ensures that the sample remains optimally hydrated. Complementing this, a compact humidifier, containing deionized (DI) water was positioned in the chamber to create a moisture environment. Post each measurement iteration of a specimen slice, an additional 20 µL of PBS is methodically dispensed onto the sample, ensuring consistent hydration conditions throughout the measurements. Due to the lower concentration of collagen and GAGs in the engineered constructs compared to native cartilage, we extended the integration time to 45 seconds. Optimal alignment of the setup was ensured before each measurement for the best signal-to-noise ratio and spectrum quality.

### Regression of purified molecular constituents

Pure components spectra of the TE constructs constituents, including collagen II (Sigma Aldrich), sulfated glycosaminoglycan (GAG) (Sigma Aldrich) and agarose (Sigma Aldrich) were measured with a confocal Raman microscope with an integration time of 1 s, employing the same spectrometer used for the RSPT setup to ensure the transferability of the spectra for the further data analysis. The spectra of water used in the multivariate data analysis were taken from a previously measured data set.^51^

## Acknowledgements

We thank the National Centre for the Replacement, Refinement and Reduction of Animals in Research (NC3Rs) for funding this work (NC/C018202/1to M.A.B.H and M.S.B). This work was supported by GlaxoSmithKline and Galvani Bioelectronics, with co-funding from the Engineering and Physical Sciences Research Council (EPSRC). M.H.Y. acknowledges funding from the China Scholarship Council (CSC). E.G. is grateful for support from the EPSRC (EP/V04723X/1). We like to thank Tianbai Wang for assisting fabrication of TE cartilage constructs.

## Contributions

E.S. and A.R.W performed the experiments, interpreted the data, generated the figures, and wrote the manuscript. D.M., M.J., and M.H.Y. performed experiments, contributed to scientific discussion, and data analysis. E.S., D.M., and S.V.P. performed experimental work and contributed to scientific discussion. E.G. and M.B.A. contributed with scientific discussion, and data interpretation. M.S.B. and M.A.B.H designed the study, interpreted the data and wrote the manuscript.

