## Supplementary information for "Label-free 3-D molecular imaging of living tissues using Raman Spectral Projection Tomography"

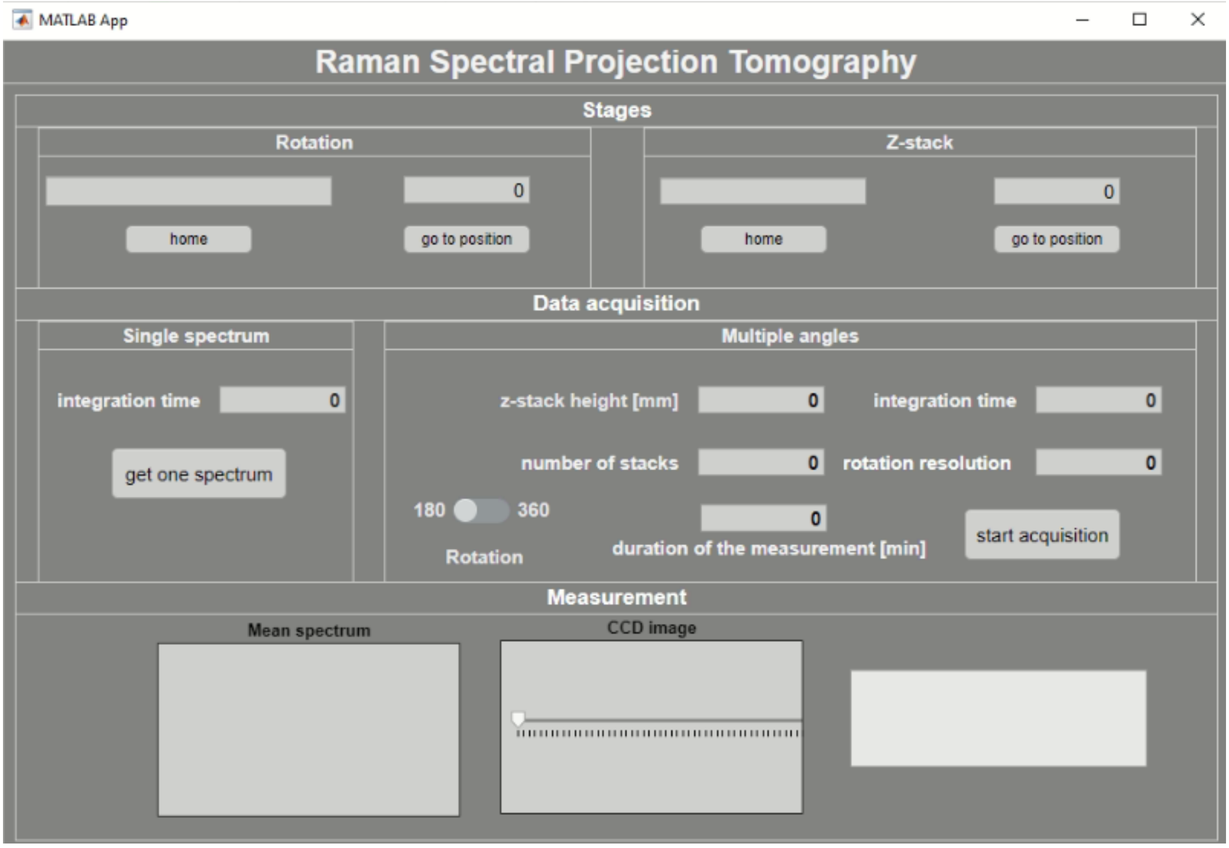


**Figure S1** Graphical User Interface (GUI) of the RSPT system software. The software facilitates automated rotation and linear translation of the stage. The system's capabilities include the ability to record a single Raman spectrum and automatically capture a comprehensive tomographic dataset. Users can adjust parameters such as the z-step height, rotational resolution, slice count, and integration duration within the software. Throughout the process, the software calculates and presents the total duration of the measurement. Users can actively observe and verify the accuracy and integrity of the ongoing measurement through a live display of the average spectrum and the unprocessed CCD image.

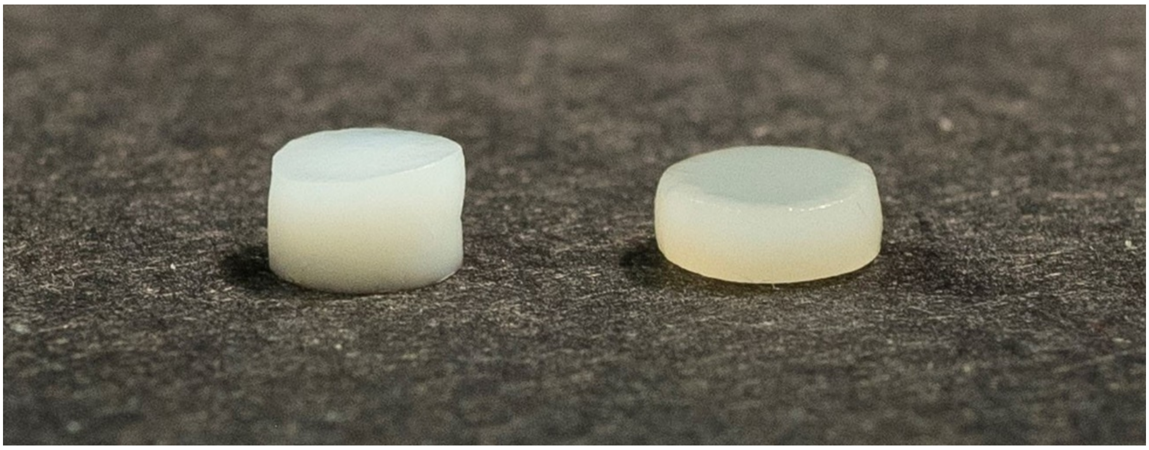

**Figure S2** Photographical comparison of native cartilage and tissue engineered cartilage. Left sample: articular cartilage explants procured from the medial and lateral condyles of 3-6 month old bovine calves. Right sample: tissue engineered cartilage after 42 days of growth.

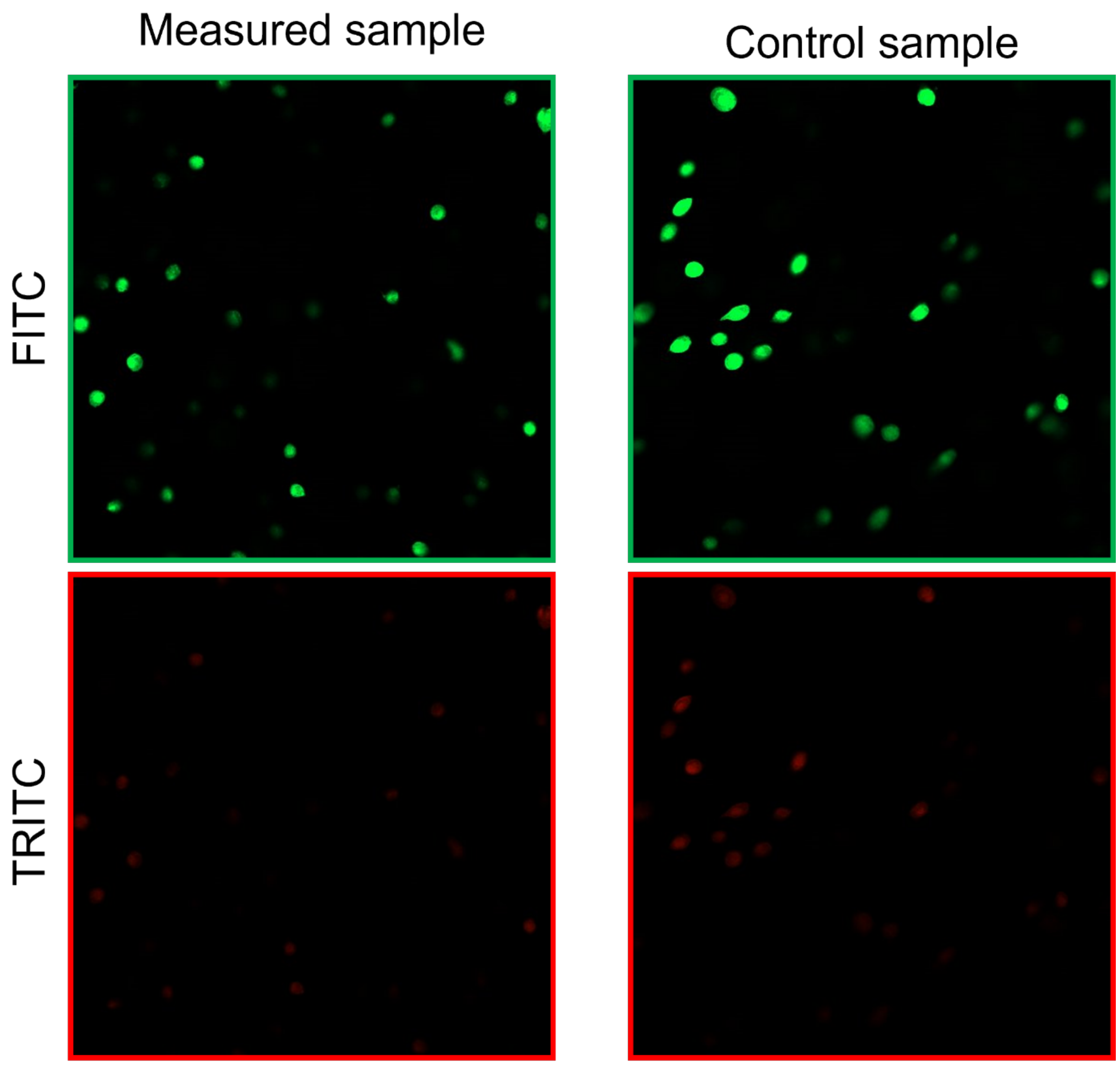


**Figure S3** Fluorescence imaging shows the outcomes of a Live/Dead assay conducted on living chondrocytes held within a hydrogel. The imaging was conducted using FITC (green) and TRITC (red) channels. Living cells emit green fluorescence, whereas deceased cells display red fluorescence. The image on the left depicts a sample exposed to high laser power intensity (2W) during Raman Spectral Projection Tomography measurement. The percentage of live cells in the laser-exposed sample was 73%.  On the right is a control group sample that was extracted from the petri dish during the measurement, offering a baseline view of cell viability. The percentage of live cells in the laser-exposed sample was 62%.

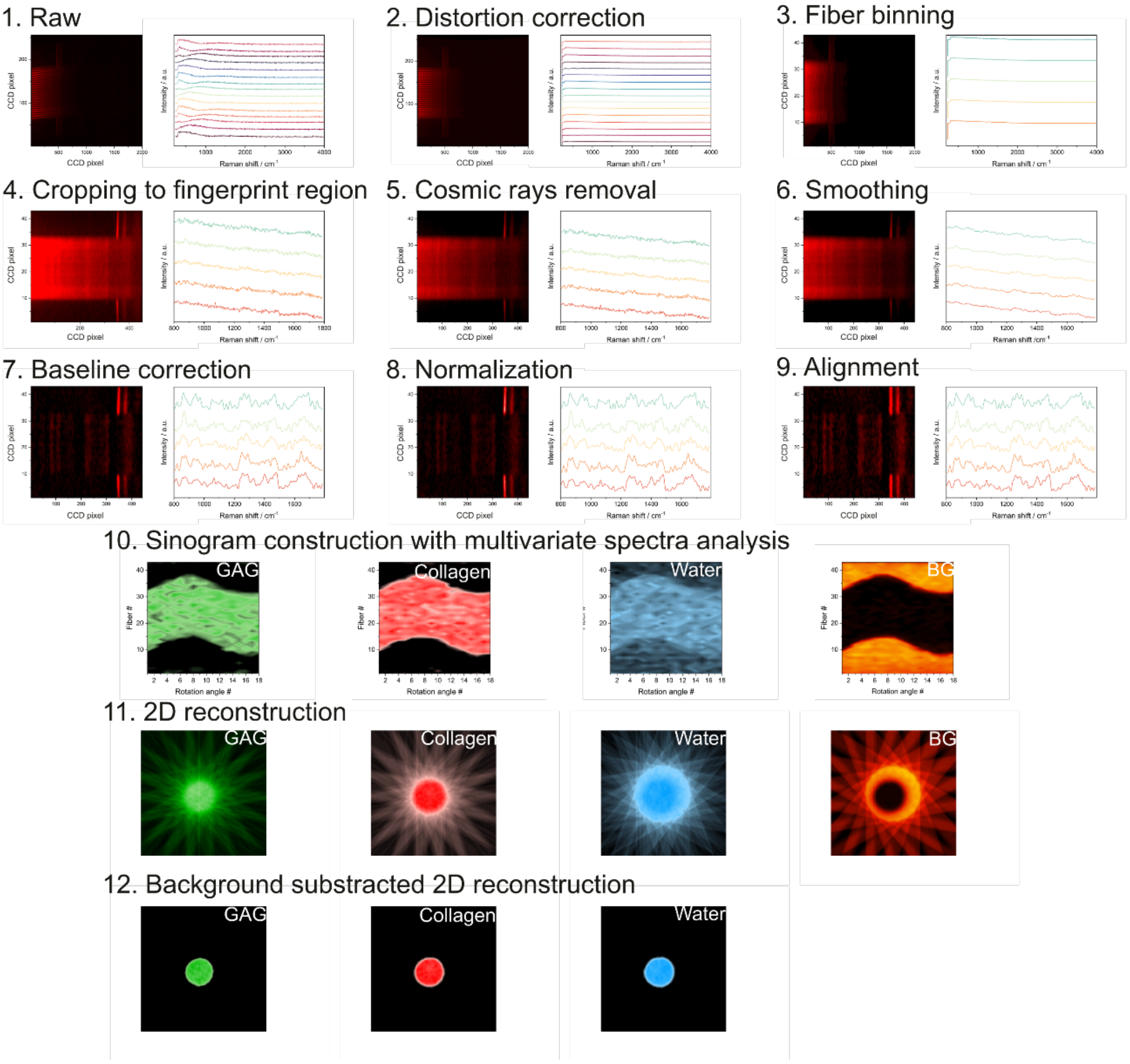


**Figure S4** RSPT Reconstruction Steps for native cartilage analysis showcasing CCD Images and corresponding Raman spectra. 1. Raw: CCD image of the cartilage sample accompanied by 10 representative raw spectra from sample region.2. Distortion correction: processed CCD image post polynomial function-based distortion correction. 3. Fiber binning: summation over every 5 rows is executed, resulting in 43 fibers. 4. Cropping to fingerprint region: spectra confined to the fingerprint region (800 $cm^{-1}$ to 1800 $cm^{-1}$) 5. Cosmic Ray removal: CCD image and spectra post removal of cosmic ray interference. 6. Smoothing: spectra subjected to a Gaussian filter for smoothing. 7. Baseline correction: Raman spectra post baseline correction via 3rd order polynomial fitting. 8. Normalization: CCD image alongside vector normalized spectra. 9. Alignment: Spectra corrected in alignment using prior measurements from a standard resin block to additionally correct for the image distortion of the CCD. 10. Sinogram construction: Constructed sinograms representing glycosaminoglycan (GAG), collagen, water, and background, derived from multivariate data analysis. 11. 2D Reconstructions: Back projected 2D reconstructions derived from the sinograms. 12. Background-subtracted 2D reconstruction: finalized 2D reconstructions after subtracting the background, leveraging the isolated background reconstruction.

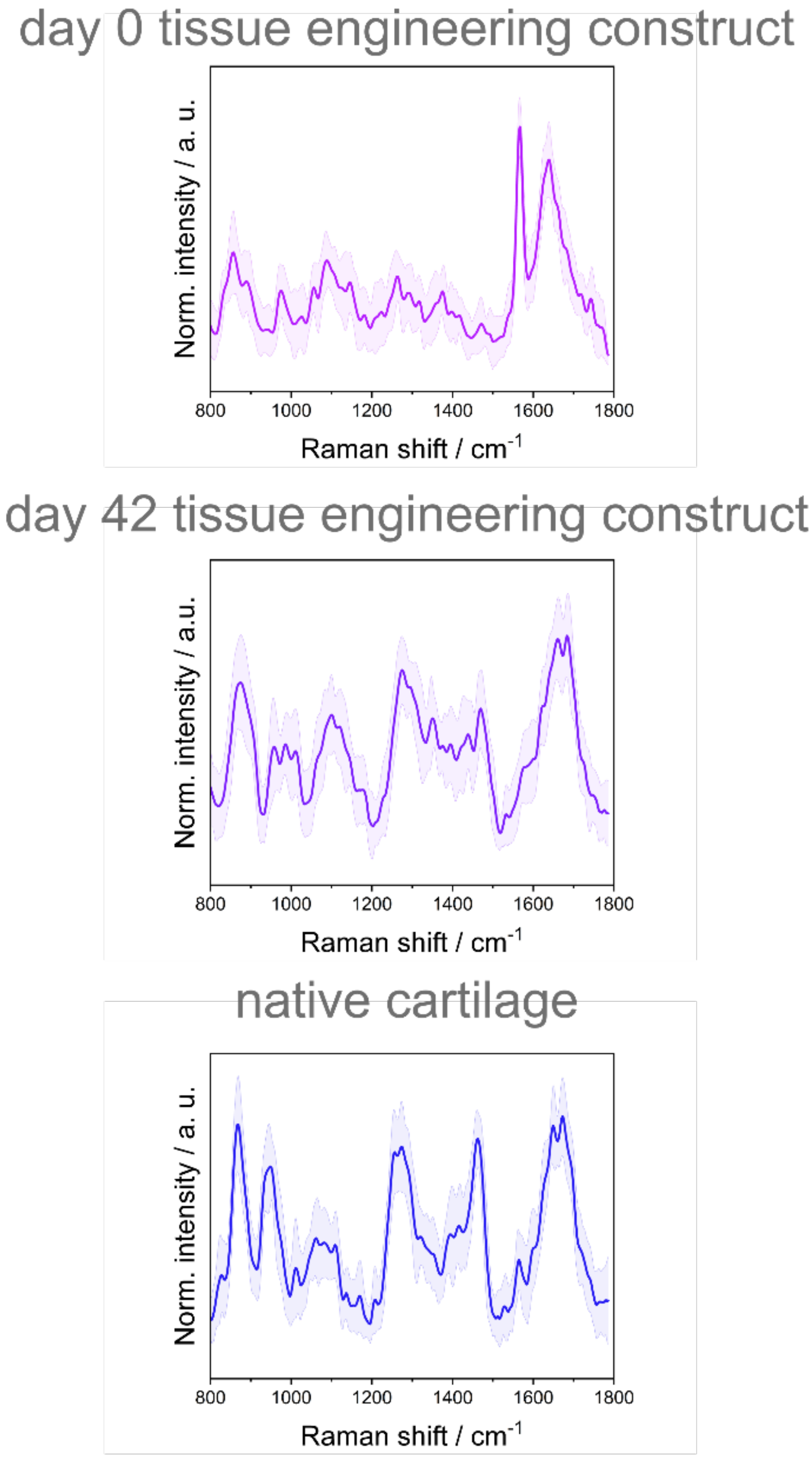


**Figure S5** Normalized mean spectra ± 1 standard deviation of tissue-engineered constructs at day 0 and day 42 and native cartilage measured with Raman Spectral Projection Tomography. Notice that for very week signals (e.g., day 0) there are minor artifacts near 1580 $cm^{-1}$ originating from the bandpass filter due to direct transmission geometry with a high intensity laser.
